# Expression profiles of hypoxia-related genes of cancers originating from anatomically similar locations using TCGA database analysis

**DOI:** 10.1101/2022.11.23.517647

**Authors:** Hye Lim Bae, Kyeonghun Jeong, Suna Yang, Hyeji Jun, Kwangsoo Kim, Young Jun Chai

## Abstract

Hypoxia is a well-recognized characteristic of the tumor microenvironment of solid cancers. This study aimed to analyze hypoxia-related genes shared by groups based on tumor location. Nine hypoxia-related pathways from the Kyoto Encyclopedia of Genes and Genomes database or the Reactome database were selected, and 850 hypoxia-related genes were analyzed. Based on their anatomical locations, 14 tumor types were categorized into the following six groups. The group-specific genetic risk score was classified as high or low risk based on mRNA expression, and survival outcomes were evaluated. The risk scores in the Female reproductive group and Lung group were internally and externally validated. In the Female reproductive group, CDKN2A, FN1 and ITGA5, were identified as hub genes associated with poor prognosis, while IL2RB and LEF1 were associated with favorable prognosis. In the Lung group, ITGB1 and LDHA were associated with poor prognosis, and GLS2 was associated with favorable prognosis. Functional enrichment analysis showed that the Female reproductive group was enriched in terms related to cilia and skin, while the Lung group was enriched in terms related to cytokines and defense. This analysis may lead to better understanding of the mechanisms of cancer progression and facilitate establishing new biomarkers for prognosis prediction.

## Introduction

Hypoxia is a common feature of the tumor microenvironment of malignant solid tumors that promotes invasive and metastatic tumor behaviors, and activates expression of various hypoxia-related genes such as hypoxia-inducible factor (1). Hypoxic foci are formed when cancer cells’ metabolic requirement surpasses the intravascular oxygen available (2). By inducing epithelial-to-mesenchymal transition, hypoxic microenvironments are associated with poor outcomes and reduced survival (3). Several hypoxia-related genes are biomarkers for the prognosis prediction of common malignancies such as breast, colorectal, gastric, and thyroid cancers (4,5).

Cancers originating in anatomically close organs may have similar genetic profiles. For example, the mutational profiles of distal colon and rectal cancer are similar although the two cancers display distinct clinical behaviors (6). Likewise, esophageal and gastric adenocarcinomas have similar mutational rates in the *APC, KRAS, PTEN*, and *SMAD4* genes (7). Hypoxia is commonly involved in tumor progression, and cancers originating in organs situated close to each other may have similar expression profiles of hypoxia-related genes.

The Cancer Genome Atlas (TCGA) database provides multiplatform genomic data of more than 20 types of carcinomas (8). It includes data about microsatellite instability, DNA sequencing, miRNA sequencing, protein expression, mRNA sequencing, DNA methylation, copy number variation, clinical information, and clinical images. TCGA project produced genomic data under standardized and controlled conditions, making it an ideal platform for pan-cancer analyses.

In this study, we hypothesized that tumors arising from the same organ or organs which are connected to each other share hypoxia-related genes. Thus, using TCGA database, we categorized 14 cancers into six groups according to their anatomical location and investigated whether cancers in the same group shared hypoxia-related genes.

## Materials and methods

### 2.1. Selection of hypoxia-related genes

We selected six hypoxia-related pathways from the Kyoto Encyclopedia of Genes (KEGG) (9,10) and three hypoxia-related pathways from the Reactome database [**Table 1**]. From the pathways, 850 genes involved in hypoxia-related pathways were selected and defined as hypoxia-related genes [**S1 Table**].

**Table 1.** Hypoxia-related Pathway list.

| Database | Pathways |
| --- | --- |
| KEGG | Pathways in cancer |
|  | Central carbon metabolism in cancer |
|  | PI3K-Akt signaling pathway |
|  | HIF-1 signaling pathway |
|  | VEGF signaling pathway |
|  | TNF signaling pathway |
| REACTOME | Hexose transport |
|  | Signaling by NOTCH1 |
|  | MAPK targets/ Nuclear events mediated by MAP kinases |

KEGG, Kyoto Encyclopedia of Genes

Fourteen tumor types in TCGA were categorized into six groups based on their anatomical locations as shown in table 2. The Liver group comprised liver hepatocellular carcinoma and cholangiocarcinoma, the Upper gastrointestinal (GI) group comprised esophageal carcinoma esophageal carcinoma and stomach adenocarcinoma, the Lower GI group comprised colon adenocarcinoma and rectum adenocarcinoma, the Female reproductive group comprised uterine corpus endometrial carcinoma and cervical squamous cell carcinoma and endocervical adenocarcinoma, the Urinary group comprised bladder urothelial carcinoma; kidney renal clear cell carcinoma; kidney renal papillary cell carcinoma; and kidney chromophobe, and the Lung group comprised lung adenocarcinoma and lung squamous cell carcinoma [**Table 2**].

**Table 2.** List of TCGA Tumor Types in Each Group.

| Group | Type |
| --- | --- |
| Liver | Liver hepatocellular carcinoma (LIHC)<br>Cholangiocarcinoma (CHOL) |
| Upper gastrointestinal | Esophageal carcinoma esophageal carcinoma (ESCA)<br>Stomach adenocarcinoma (STAD) |
| Lower gastrointestinal | Colon adenocarcinoma (COAD)<br>Rectum adenocarcinoma (READ) |
| Female reproductive | Uterine corpus endometrial carcinoma (UCEC)<br>Cervical squamous cell carcinoma and endocervical<br>adenocarcinoma (CESC) |
| Urinary | Bladder urothelial carcinoma (BLCA)<br>Kidney renal clear cell carcinoma (KIRC)<br>Kidney renal papillary cell carcinoma (KIRP)<br>Kidney chromophobe (KICH) |
| Lung | Lung adenocarcinoma (LUAD) |
|  | Lung squamous cell carcinoma (LUSC) |

TCGA, The Cancer Genome Atlas

The TCGA pan-cancer RNA-seq data and the TCGA-Clinical data resource outcome were downloaded from the PanCanAtlas publications page (https://gdc.cancer.gov/about-data/publications/pancanatlas). A total of 4,751 tumor samples with TCGA-CDR outcome data were used for downstream analysis.

Among the 850 hypoxia-related genes, six genes (*LOC101928143, LOC102723407, MIR1281, PLA2G4B, SLC5A10*, and *G6PC1*) were excluded from TCGA pan-cancer RNA-seq data, thus, 844 genes with RNA expression values were used in the analysis.

### 2.2. Group-specific genetic risk gene identification

We used Cox regression to calculate the group-specific genetic risk score for each group based on mRNA expression of hypoxia-related genes as the predictor variable. Elastic net penalized Cox regression analysis was conducted based on the RNA expression values of 844 hypoxia-related genes and each group’s overall survival data. Genes with Cox regression model coefficients that were not equal to 0 were defined as group-specific genetic risk genes. The group-specific genetic risk score, Rg, was calculated using the following equation:

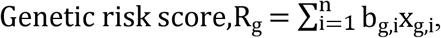

where *g* represents group classification according to anatomical location, *i* represents the number of genes with coefficients not equal to 0, *b* represents the elastic net Cox regression coefficient value of each gene, and *x* represents the RNA expression value of each gene.

*Glmnet* R package was employed for elastic net Cox regression analysis.

### 2.3. Survival analysis and internal validation

Group samples were classified as ‘high-risk’ or ‘low-risk’ based on the median value of the group-specific genetic risk score. Survival analysis was conducted using Kaplan-Meier survival curves. Log-rank tests were performed to compare survival curves between high-risk and low-risk groups.

Tumor types with log-rank test p-values between high-risk and low-risk classes below 0.05 across all tumor types within each group were defined as internally validated groups.

R package *pROC* was used for building receiver operating characteristic(ROC) curves and *survminer* was used for the Kaplan-Meier plot and log-rank test.

### 2.4. External validation

To improve the validity and generalizability, external validation was performed on the internally validated groups, using the dataset of the Gene Expression Omnibus (GEO).

Nineteen external validation datasets were obtained from the GEO database [**S3 Table]**. Log2 transformed raw expression values were used for risk score calculation. For each dataset, samples were classified as high- and low-risk by genetic risk score calculated. We calculated the gene coefficients of the group. R packages *GEOquery, Affy, Oligo* and *Limma* were used for the pre-processing of external validation datasets. To detect and remove outlier, *outlier_osd* function of the R package *survBootOutliers* was applied.

Survival analysis was performed by dividing samples of each dataset into high- and low-risk classes based on the median genetic risk score.

### 2.5. Functional enrichment analysis

Gene set enrichment analysis (GSEA) was conducted using the R package *clusterProfiler*. KEGG pathway, Gene Ontology Biological Process terms from the Molecular Signatures Database were used to perform functional annotation of differentially expressed genes (DEGs) between high- and low-risk classes within each validated group (11). The top ten enriched terms were extracted in this study.

Among the genes with an average of more than three RNA expression values, DEGs between high- and low-risk classes within each group were identified with a cutoff value at Wilcoxon rank-sum test false discovery rate < 0.01 and absolute log2 fold change > 1.

### 2.6. Gene ontology analysis

Gene ontology analysis was conducted using the R package *clusterProfiler* (12). KEGG pathway, Gene Ontology biological process terms from the Molecular Signatures Database were used to perform functional annotation of positive risk-coefficient genes within each validated group (11). The 844 hypoxia-related genes were used as a background gene set. Gene ontology terms less than a p-value of 0.5 were defined as the enriched term. The top 10 terms among enriched terms less than p-value of 0.2 were represented as dot plots.

## Results

### 3.1. Selection of hypoxia-related genes

From the hypoxia-related pathways, 850 genes involved in hypoxia-related pathways were selected and defined as hypoxia-related genes **[S1 Table]**. Among the 850 hypoxia-related genes, six genes (*LOC101928143, LOC102723407, MIR1281, PLA2G4B, SLC5A10*, and *G6PC1*) were absent from TCGA pan-cancer RNA-seq data, thus, 844 genes with RNA expression values were used in the analysis.

### 3.2. Group-specific genetic risk score identification

Group-specific genetic risk genes and coefficients are listed in Table 3 **[S2 Table]**.

**Table 3.** List of Group-specific Hypoxia Risk Genes.

| Group | Positive risk coefficient | Negative risk coefficient |
| --- | --- | --- |
| Liver | *BIRC5*, *BIRC8*, *CUL2*, *EIF4E*, *EPO*,<br>*G6PD*, *GNAI2*, *HDAC1*, *HDAC2*,<br>*HSP90AA1*, *IFNA13*, *IL8*, *LDHA*,<br>*MAPK7*, *NUP155*, *PGF*, *PPP2R5B*,<br>*RHEB*, *SLC1A5*, *SLC2A1*, *SPP1*,<br>*YWHAB* | *CCNA1*, *CNTN1*, *FLT3*, *G6PC2*,<br>*GHR*, *HES5*, *IFNA2*, *ITGB7*, *NTRK1*,<br>*PFKL*, *TNF*, *TP53*, *WNT1* |
| Upper gastrointestinal | *APH1B, SERPINE1, SLC2A3, SOCS3, TF* | *DAB2IP, MKNK2* |
| Lower gastrointestinal | *APC2, ENO3, HEYL, TIMP1, WNT10B* | *CTNNA1, MAPKAPK3, TMEM48* |
| Female reproductive | *BDKRB1, CDKN2A, FN1, ITGA5, PFKM, SLC45A3, TFRC, VEGFA, WNT3, YWHAB, YWHAG* | *CD19, IL2RB, JMJD7-PLA2G4B, LEF1, MDM2, RBPJ* |
| Urinary | *BIRC5, CCNE2, COL6A3, DVL3, EIF4EBP1, FGF5, GLI2, PPP2R2C, SLC7A5, THBS3* | *DAB2IP, ITGB7* |
| Lung | *ITGB1, LDHA* | *GLS2* |

The distribution of hypoxia genetic scores is shown in the risk score plot **[Fig 1]**. In the Female reproductive group, 232 (79%) out of the 293 patients with CESC tumor type were classified as high-risk. Risk scores were evenly distributed across tumor types in the Upper GI, Lower GI, and Lung groups.

**Fig 1.**
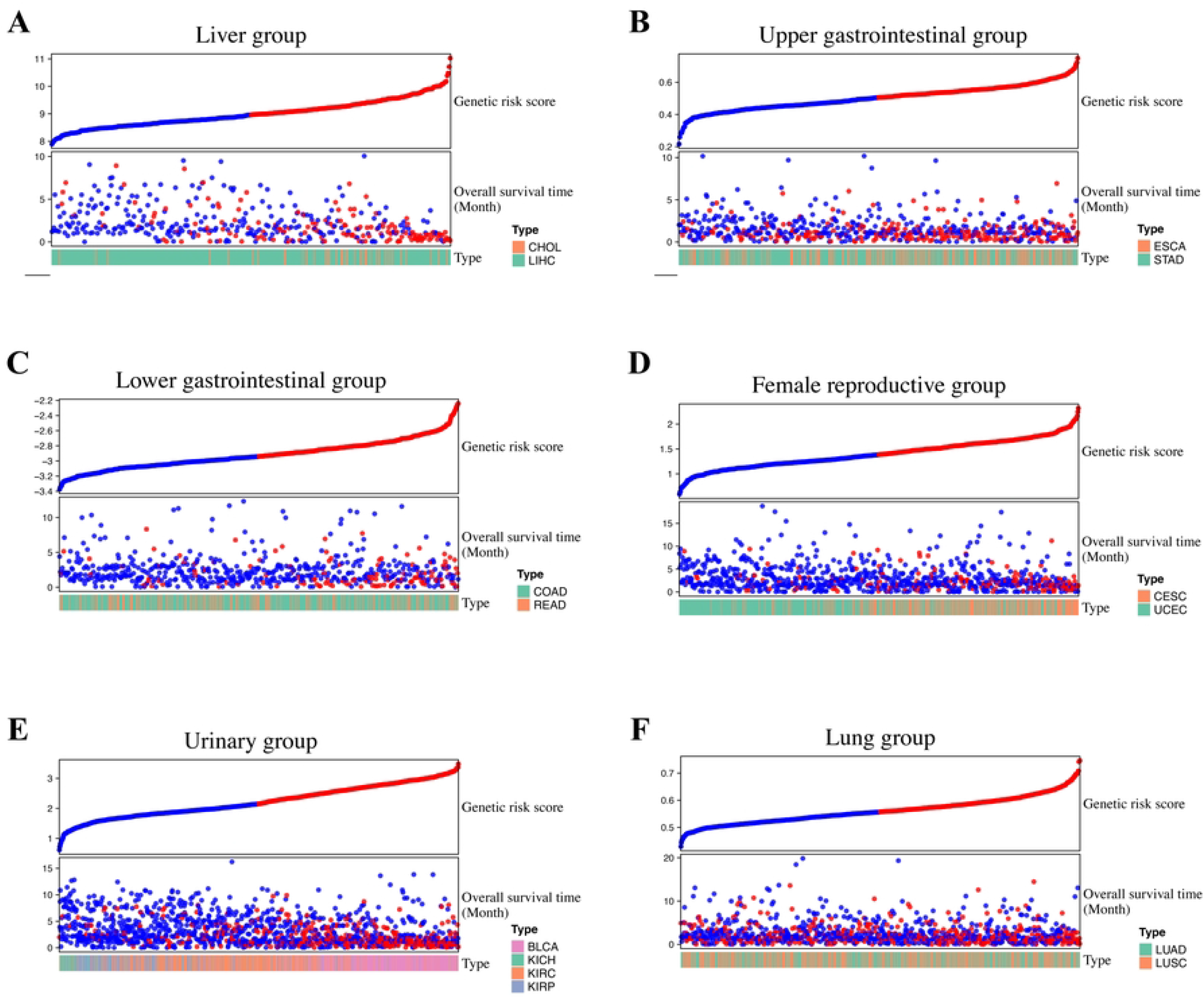
Risk score distribution plot of each group. In the Female reproductive group, 79% of patients with CESC tumor type were classified as high-risk. Risk scores were evenly distributed across tumor types in the Upper GI, Lower GI, and Lung groups. (a) Liver group (b) Upper GI group (c) Lower GI group (d) Female productive group (e) Genitourinary group (f) Lung group

### 3.3. Survival analysis and internal validation

Overall survival between high- and low-risk classes were compared using the Kaplan-Meier method and the log-rank test **[Fig 2]**. To assess risk score performance of each tumor, samples were divided into high- and low-risk classes based on the median risk score within each group, and survival was compared using the Kaplan-Meier method and log-rank test. The Female reproductive, Lung, Upper GI, and Lower GI groups were internally validated **[Fig 3]**. The log-rank test p-values comparing survival of high- and low-risk are listed in Table 4. Except for CHOL and KICH, the remaining cancer types showed significantly different results. (p-value < 0.05)

**Fig 2.**
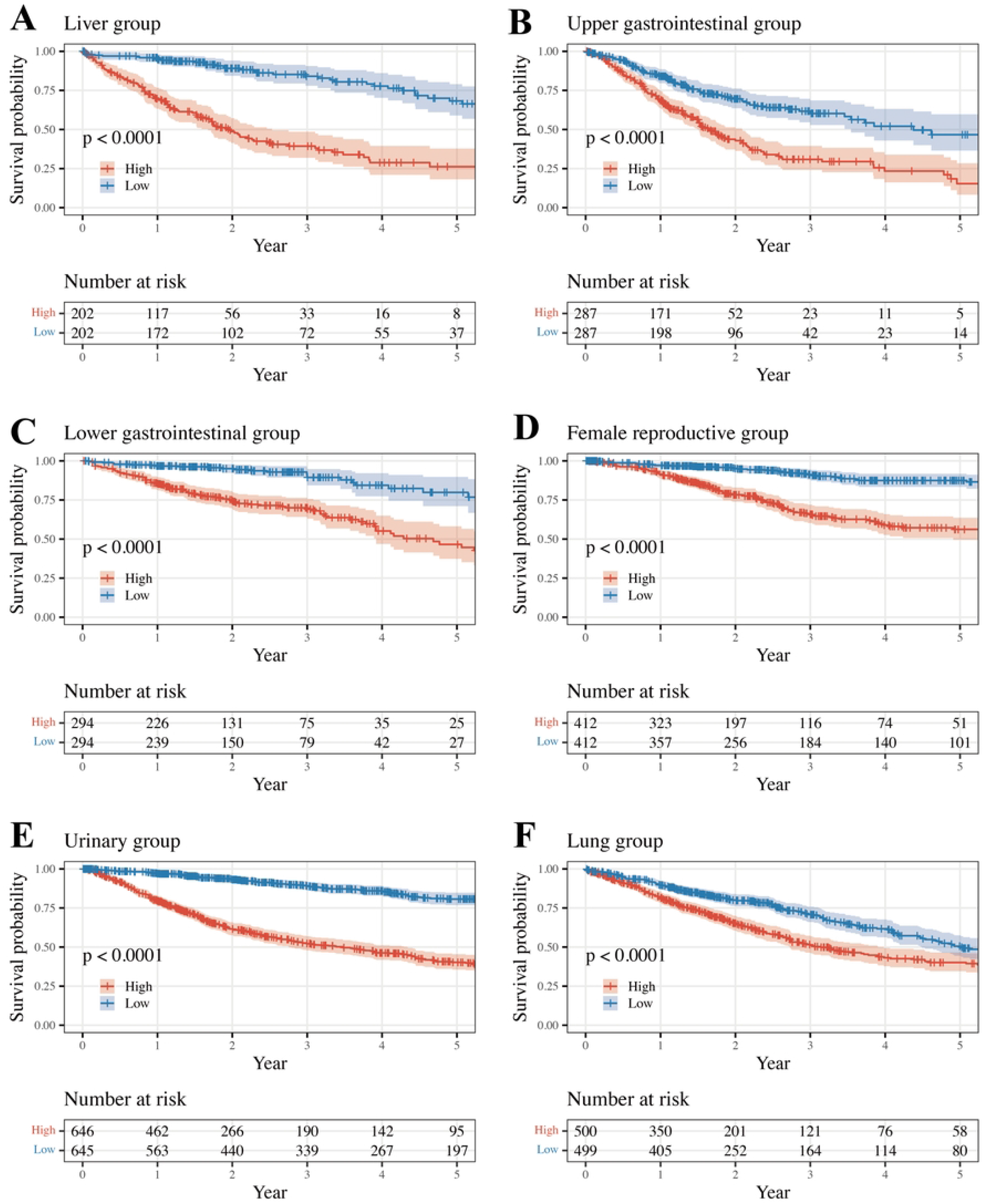
Comparison of Kaplan-Meier survival curves between high- and low-risk classes. (a) OS in Liver group (b) OS in Upper GI group (c) OS in Lower GI group (d) OS in Female productive group (e) OS in Genitourinary group (f) OS in Lung group, OS overall survival; GI, gastrointestinal

**Fig 3.**
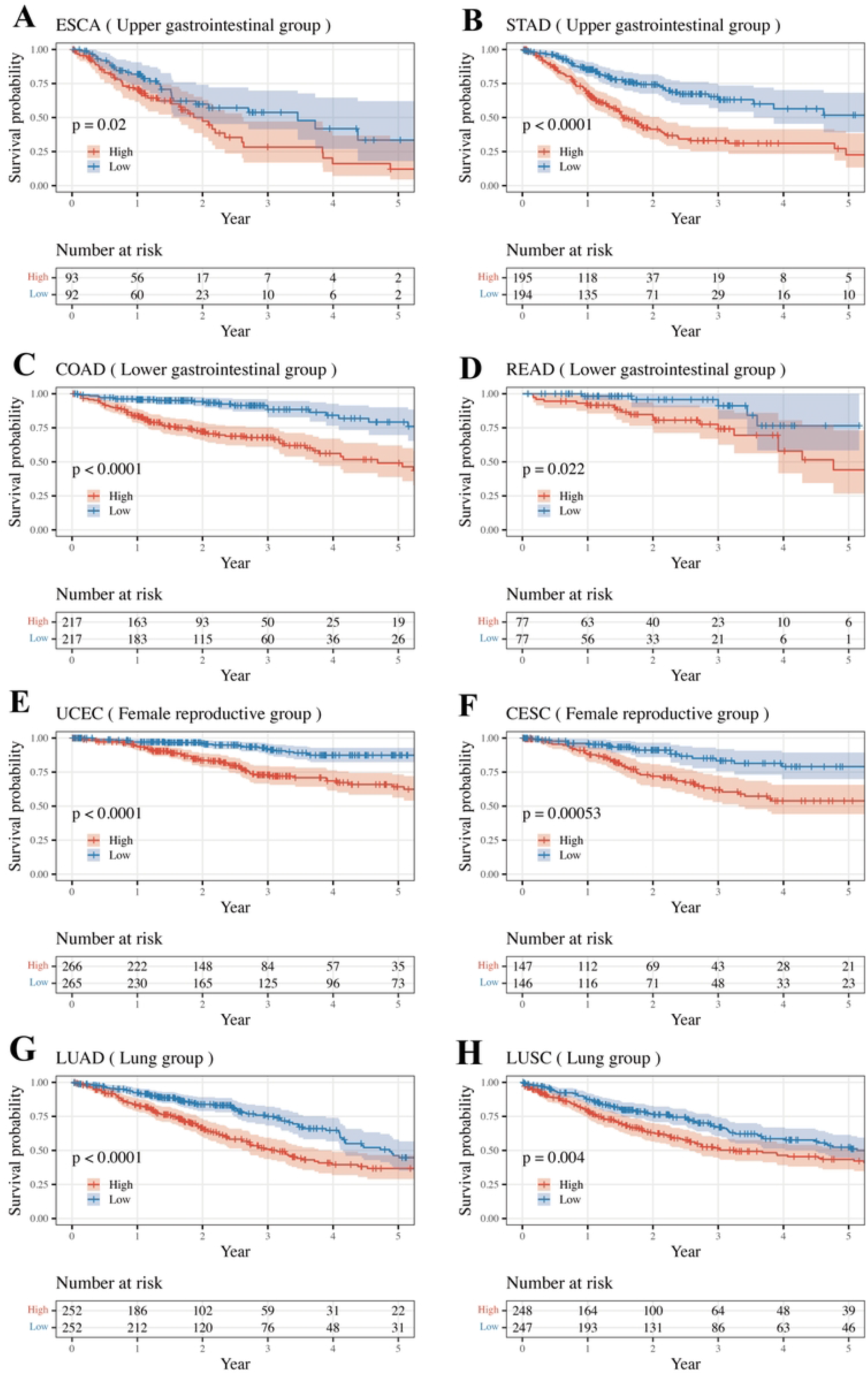
Comparison of Kaplan-Meier survival curves between high- and low-risk classes based on the median risk score. The Female reproductive, Lung, Upper GI, and Lower GI groups were internally validated. (a) OS in ESCA (b) OS in STAD (c) OS in COAD (d) OS in READ (e) OS in UCEC (f) OS in CESC (g) OS in LUAD (h) OS in LUSC, OS overall survival;

**Table 4.** Group log-rank test results and TCGA internal validation results.

| Group | Group-specific log rank p-value | Type | Type-specific log rank p-value | Internal validated |
| --- | --- | --- | --- | --- |
| Liver | $5.12 \times 10^{-18}$ | LIHC* | $2.92 \times 10^{-17}$ | FALSE |
| | | CHOL | $6.17 \times 10^{-2}$ | |
| Upper gastrointestinal | $1.39 \times 10^{-9}$ | ESCA | $1.99 \times 10^{-2}$ | TRUE |
| | | STAD* | $7.88 \times 10^{-8}$ | |
| Lower gastrointestinal | $8.02 \times 10^{-11}$ | COAD* | $5.20 \times 10^{-9}$ | TRUE |
| | | READ | $2.19 \times 10^{-2}$ | |
| Female reproductive | $9.10 \times 10^{-16}$ | UCEC* | $1.51 \times 10^{-7}$ | TRUE |
| | | CESC | $5.32 \times 10^{-4}$ | |
| Urinary | $9.86 \times 10^{-47}$ | BLCA* | $3.46 \times 10^{-5}$ | FALSE |
| | | KIRC* | $3.75 \times 10^{-13}$ | |
| | | KIRP* | $7.53 \times 10^{-5}$ | |
| | | KICH | $8.29 \times 10^{-2}$ | |
| Lung | $1.51 \times 10^{-6}$ | LUAD* | $7.06 \times 10^{-5}$ | TRUE |
| | | LUSC | $3.95 \times 10^{-3}$ | |

\*p-value < 0.0001

### 3.4. External validation result of internally validated groups

Risk scores were calculated using the group-specific genetic risk score gene coefficients calculated from TCGA samples **[Table 5]**.

**Table 5.** Datasets used for external validation and log rank test results.

| Group | Matched TCGA Type | GEO Accession number |
| --- | --- | --- |
| Upper gastrointestinal | STAD | GSE15459* |
|  | ESCA | GSE72873 |
| Lower gastrointestinal | COAD<br>READ | GSE41258, GSE17538, GSE72970,<br>GSE17537, GSE17536 |
| Female reproductive | CESC | GSE52903* |
|  | UCEC | GSE119041* |
| Urinary | BLCA | GSE31684, GSE13507, GSE19423 |
|  | KIRC | GSE29609 |
| Lung | LUAD/SC | GSE11969*, GSE37745 |
|  | LUAD | GSE31210*, GSE30219, GSE50081,<br>GSE29014 |

\*p-value < 0.05

Significant differences in overall survival curves between the high- and low-risk classes were seen in the Lung group, GSE11969 (13,14) (matching TCGA type: LUAD and LUSC) and GSE31210 (15) (matching TCGA type: LUAD), and the Female reproductive group, GSE119041 (16) (matching TCGA type: UCEC), and GSE52903 (17) (matching TCGA type: CESC) **[Fig 4, S3 Table]**. Accordingly, the Female reproductive and Lung groups were internally and externally validated.

**Fig 4.**
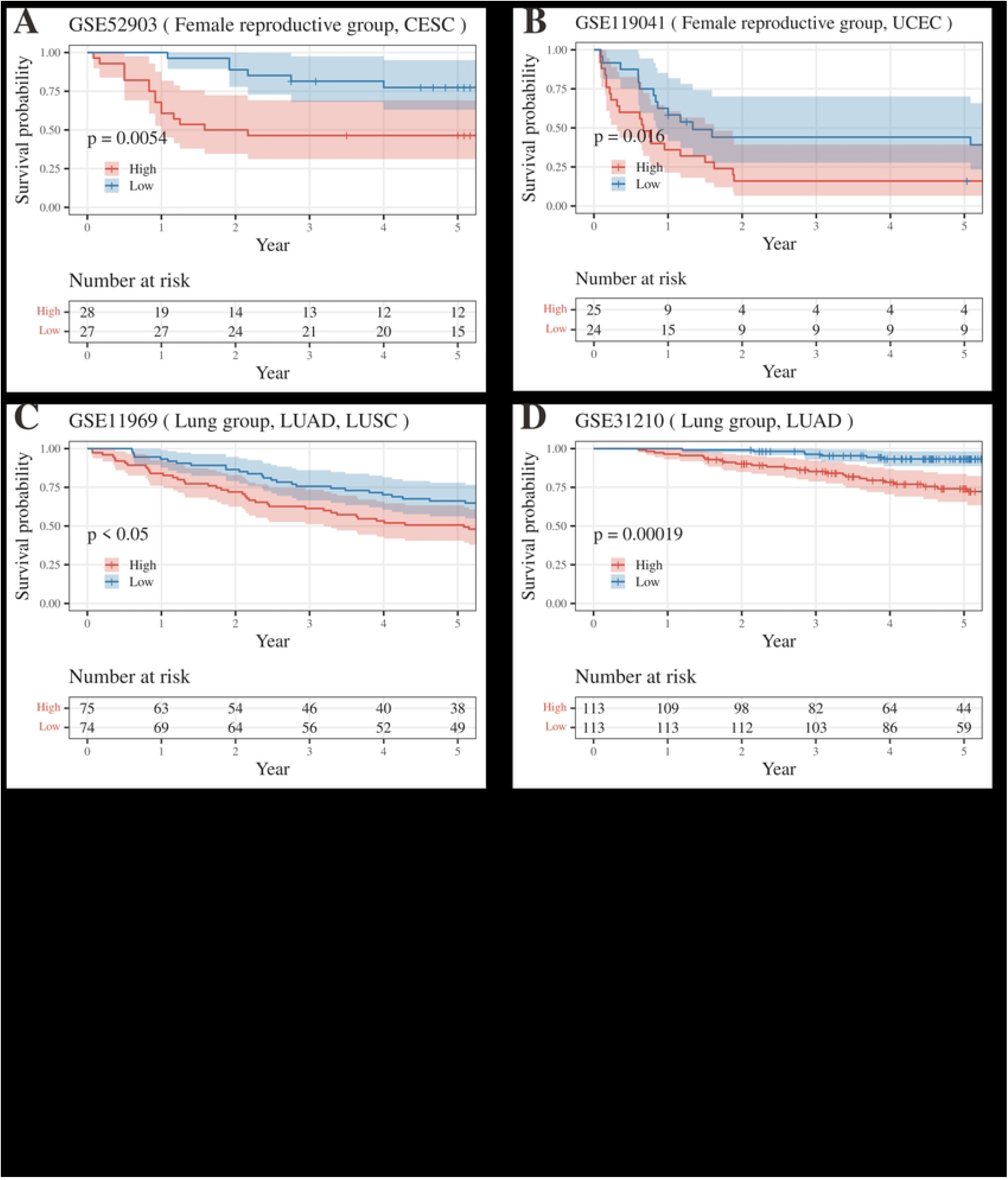
Kaplan-Meier plot of each external validation dataset. The Female reproductive and Lung groups were externally validated. **(a)** OS in GSE52903 **(b)** OS in GSE119041 **(c)** OS in GSE11969 **(d)** OS in GSE31210, OS overall survival;

### 3.5. Functional enrichment analysis

Functional enrichment analysis was performed to identify differences in the pathway between the high- and low-risk classes for the internally and externally validated groups.

GSEA shows that 1,638 DEGs between high- and low-risk classes in the Female productive group were enriched in terms related to cilia (‘cilium movement’ and ‘cilium organization’) and terms related to skin (‘skin development’ and ‘epidermis development’). **[Fig 5; S4 and S5 Tables]**

**Fig 5.**
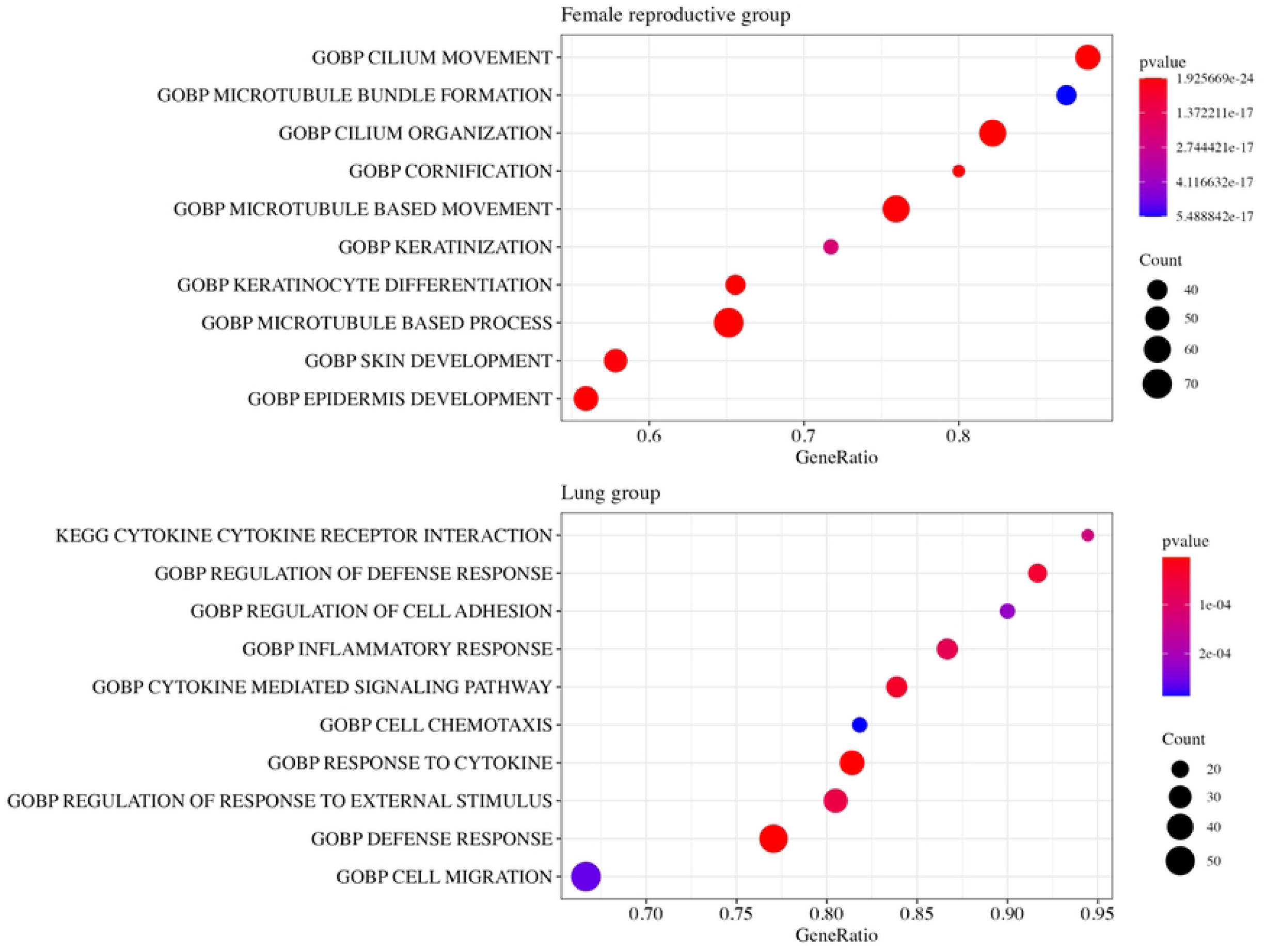
Top 10 enriched terms in the Female productive group and the Lung groups. The Female productive group and the Lung groups are internally and externally validated groups. DEGs in the Female productive group were enriched in terms related to cilia, while DEGs in the lung group were enriched in terms related to cytokines.

In the Lung group, 505 DEGs were enriched in terms related to cytokines (‘cytokine’, ‘cytokine receptor interaction’, ‘response to cytokine’, and ‘cytokine mediated signaling pathway’) and defense (‘regulation of defense response’ and ‘defense response’). **[Fig 5; S6 and S7 Tables]**

### 3.6. Gene ontology analysis

The gene ontology (GO) analysis was performed to identify the function of positive risk-coefficient genes.

GO analysis shows that 11 positive risk-coefficient genes (*BDKRB1, CDKN2A, FN1, ITGA5, PFKM, SLC45A3, TFRC, VEGFA, WNT3, YWHAB, YWHAG*) in the Female productive group were enriched in terms related to axon regulation, such as POSITIVE REGULATION OF AXONOGENESIS, AXON EXTENSION, and POSITIVE AXON REGULATION OF AXON EXTENSION. **[Fig 6**, **S8 Table**]

**Fig 6.**
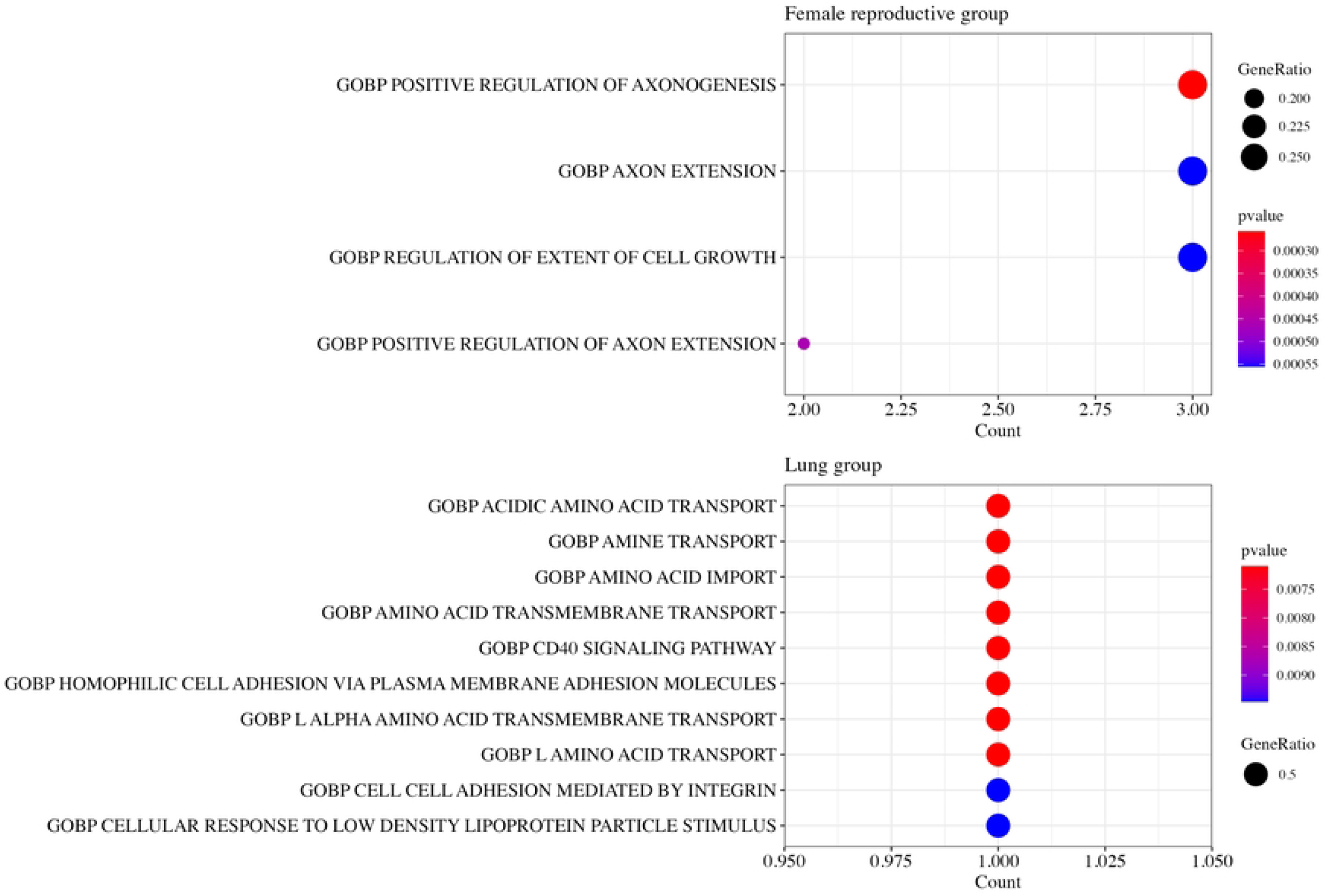
Top 10 enriched terms in the Female productive group and the Lung groups. The Female productive group and the Lung groups are internally and externally validated groups. The positive risk-coefficient genes in the Female productive group were enriched in terms related to axon regulation, while those of the lung group were related to amino acid transport.

Two positive risk-coefficient genes (*ITGB1, LDHA*) in the lung group were related to amino acid transport. Terms such as ACIDIC AMINO ACID TRANSPORT, AMINE TRANSPORT, AMINO ACID IMPORT, etc. were enriched. [**S9 Table**]

## Discussion

Hypoxia induces metabolic and molecular changes in the majority of malignant tumors. The association between the genomic characteristics of hypoxia and aggressive tumor cell phenotypes is well-established (18). However, few studies have investigated the genetic profile of hypoxia-related genes of cancers originating from anatomically close organs and the genomic profiles from large scale. We performed analysis of hypoxia-related genes by categorizing 14 cancers into six groups originating from the same or adjacent organs based on TCGA data and validated the results using external dataset. We found that the prognosis of Female reproductive and Lung groups differed significantly between low-risk and high-risk group, affected by hypoxia-related genes.

CESC and UCEC are representative gynecological cancers. Although these two cancer types have different clinical features, origins, and prognosis, studies have demonstrated gynecologic cancers share abnormally-expressed genes (19,20). Two hub genes (*PAMR1* and *SLC24A3*) are potential shared biomarkers for both CESC and UCEC (20). The expression of *PAMR1* and *SLC24A3* in cancer tissue is downregulated significantly compared with normal tissue. *PAMR1* influences epithelial-to-mesenchymal transition by inhibiting the proliferation, migration, and invasion of cancer cells (21). *SLC24A3* (also known as *NCKX3*) is involved in the transport of calcium across the cell. Its expression is abundant within the human endometrium at the mRNA and protein levels, especially during menstruation, and has a role in the reproductive cycle (22). Another study found that *MAL* overexpression may predict poor prognosis in CESC and UCES (23). One study reported that *MAL* expression increased in chemo-resistant cancers, and is associated with short overall survival (24). Expression of *ACTA1, MYH7*, and *MYBPC1* may be a potential promotor of gynecological cancer initiation or progression (19). These genes are regulators of actin and myosin and, as such, changes in actin bundling proteins caused by gene alterations could be correlated with cancer initiation or progression.

In agreement with published studies, the current study identified overlapping molecular finding in gynecological cancer, focusing on hypoxia gene. We identified 11 positive risk coefficient genes and seven negative risk coefficient genes. Among them, *CDKN2A* (also known as *P16* gene), *FN1*, and *ITGA5* were identified as tumor markers which predict poor prognosis. *CDKN2A* encodes tumor suppressor protein or tumor immunity (25). *CDKN2A* methylation has been reported in association with poor prognosis in ovarian cancer (26). The *FN1* gene is a glycoprotein involved in cell proliferation and migration. The expression of *FN1* is a very poor prognosis marker for various cancer types, including gastric, colorectal, and thyroid cancers (21,27). Other studies report that expression of *ITGA5* is increased in breast, stomach, and ovarian cancers, compared with normal tissue (28–30). *ITGA5* is a member of the integrin alpha chain family and combines with *ITGB1* to form integrin α5β1, which has been demonstrated to engage in tumor cell adherence (31). Progression of gynecological cancer may be promoted by micro-environmental changes in tumor immunity, adherence, and migration caused by alteration of hypoxia-related genes.

We also found that upregulation of certain genes was associated with better prognosis. In the current study, *IL2RB* and *LEF1* were associated with good prognosis in the Female reproductive group. In contrast to the current findings, high expression of *IL2RB* and *LEF1* has been correlated with poor prognosis (32). *IL2RB*, as a T-cell-mediated immune system regulating gene, was reported to increase cytotoxic lymphocytes, T-cells, and natural killer cells, leading to immune invasion and tumorigenesis (33). *LEF1* is essential for T- and B-cell differentiation and its transcription factors are required for self-renewal of leukemic stem cells (34). Thus, LEF1 is consistently associated with T-cell tumors in the literature (35).

The GSEA data illustrated that the cilia and epidermis have a vital function in gynecological tumorigenesis. The human endometrium comprises abundant motile cilia, and ciliary defect may have a role in the early stage of tumor development. When the motile cilia decrease, oxidative stress in epithelial cells is exacerbated, which can lead to precursor cancer (36). The exocervix and vagina are lined with squamous epithelium that form the surface of the skin and hollow organs. If these cells repeatedly suffer from external insult (e.g., repeated infection by human papillomaviruses), cell cycle regulatory tumor suppressor proteins, such as *p53* and *pRB*, are inactivated allowing epidermal cells in squamous cell carcinoma to abnormally proliferate and dedifferentiate (37). In gynecological cancers, the functional pathways of epithelial cells are the most important mechanism of cancer progression.

GO analysis showed that axon control is one of the influential factors in the micro-environment of Female reproductive cancer. Recently a study denoted the connection between neuro activity and cancer cell growth (38). The release of neurotransmitters stimulate around cancer cell and its stromal cell. The process is also facilitated by growth factor of cancer cells, leading to perineural invasion. Other research also suggested that neural activity such as axonogenesis is also observed in gastric and colon cancer (39,40). Our finding also shows the similar trend with previous studies, which is nerve is a crucial role in tumor infiltration around cells.

Lung adenocarcinoma and lung squamous cell carcinoma are major cancers which have been investigated by many studies. Although they originate from different cells and have different molecular profiles, some studies have demonstrated common gene pathways. Six genes (*PGK1, ENO2, GPI, PEKP, ALDOA* and *ANGPTL4*) were reported as hypoxia-related genes in lung cancer (41). These genes function as regulators of oxygen-dependent molecular pathways, leading to increased anaerobic glycolysis. *TTF1, KRT7, SOX2, P63* and *KRT5* are biomarkers for poor prognosis of lung adenocarcinoma and lung squamous cell carcinoma (42). Among these genes, *TTF1* was demonstrated to contribute to the maintenance of the function of terminal respiratory unit cells and is used in immunohistochemistry differential diagnosis of lung cancer (43). *SOX2*, as a stem cell transcription factor, regulates human somatic cells to pluripotent stem cells (44). One study revealed that overexpression of *SOX2* amplifies the 3q gene, which is the most common genomic mutation in lung cancer (45).

In this study, *ITGB1* and *LDHA* were poor prognosis markers and *GLS2* was an indolent marker. *ITGB1* has been reported to regulate cancer migration, invasion, and metastasis. Previous studies have shown that knockdown of *ITGB1* reduces breast cancer and colorectal cancer (46,47). *LDHA* is an enzyme gene involved in creating an acidic microenvironment by effecting the pyruvate cycle. When *LDHA* is overexpressed, epithelial-to-mesenchymal transition is overactivated, which is associated with poor prognosis (48). In contrast *GLS-2* is has recently been identified as a key gene in the suppression of cancer metastasis via regulating glutamine metabolism (49). It binds to *Rac1-GDP* by inhibiting Rac1 activity, which eventually activates the *p53* tumor suppression gene. *GLS-2* is recently identified as a key gene in the suppression of cancer metastasis. The modification of proteins and other anti-oncogenes by alteration of hypoxia-related genes may determine cancer prognosis.

Functional enrichment analysis indicated that the molecular mechanisms related to cytokine and defense were enriched in lung cancer. Despite controversy that immune proteins alone are not related to risk cancer, cytokine expression in lung cancer has been described in the recent research (50). Tumor cells yield immunosuppressive cytokines, which is able to avoid host immune attack (51). The impaired anticancer defense system can also cause avoiding cancer cells from immune system (52). These mechanisms of immunogenicity in lung cancer may potentially have a crucial function in the regulation of cancer progression.

GO analysis revealed that change of amino acid profiles accelerate the tumor growth in lung cancer. It is known that particular amino acids facilitate the proliferation of cancer cells and potential role in regulation of microenvironment (53). Among them, tryptophan is a crucial amino acid for immune proliferation by regulating T-cells (54). The change of tryptophan can induce immune escape, leading to promotion of cancer cells. In addition, asparagine, aspartate, and glutamine are known as intracellular and extracellular amino acids which serve to assist in cell integration (55). When the amino acids are depleted, it cause impairment of protein synthesis and finally lead to unusual apoptosis. All these metabolism shifts can induce to change microenvironment for lung cancer to grow well.

The strength of the current study was the analysis of hub genes according to the location of cancer, based on massive data mining. However, there are some limitations. First, as a retrospective study, there is inevitable selection bias. Therefore, we performed external validation with larger sample sizes to overcome the bias. Second, detailed information about the tumors such as staging, grading, and prognosis were not included in the original dataset and were thus omitted from analysis. Therefore, further prospective clinical trials and experimental studies are needed to further validate our findings.

## Conclusion

We identified common hypoxia-related genes in Female reproductive cancers as well as lung cancers and detected functional mechanisms to further elucidate the developmental process of cancer. This bioinformatics analyses expands our knowledge of cancer, and may lead to the development of better personalized treatment strategies. Extensive research related to the hypoxia genes are required to predict cancer prognosis through risk stratification.

## Supporting information

**S1 Table. Genes associated with Hypoxia-related pathway**

**(XLSX)**

**S2 Table. A list of group-specific hypoxia risk genes and coefficients**

**(XLSX)**

**S3 Table. Datasets Used for External Validation and Log Rank Test Results**

**(XLSX)**

**S4 Table. DEG analysis results in the Female reproductive group**

**(XLSX)**

**S5 Table. Gene set enrichment analysis results for DEGs in the Female reproductive group**

**(XLSX)**

**S6 Table. DEG analysis results in the Lung group**

**(XLSX)**

**S7 Table. Gene set enrichment analysis results for DEGs in the Lung group**

**(XLSX)**

**S8 Table. GO analysis results for Female reproductive group positive risk-coefficient genes**

**(XLSX)**

**S9 Table. GO analysis results for Lung group positive risk-coefficient genes**

**(XLSX)**

## Acknowledgment

We thank all members of K.K. and C.Y.J. labs for beneficial discussions. Availability of data and materials from TCGA dataset (https://tcga-data.nci.nih.gov/tcga/).

## Author Contributions

**Conceptualization:** Kwangsoo Kim, Young Jun Chai

**Data curation:** Kyeonghun Jeong

**Formal analysis:** Hye Lim Bae, Kyeonghun Jeong

**Funding acquisition:** Young Jun Chai

**Investigation:** Kyeonghun Jeong

**Methodology:** Suna Yang, Hyeji Jun

**Project administration:** Hye Lim Bae

**Software:** Hye Lim Bae, Kyeonghun Jeong

**Resources:** Suna Yang, Hyeji Jun

**Supervision:** Kwangsoo Kim, Young Jun Chai

**Validation:** Kyeonghun Jeong

**Visualization:** Suna Yang, Hyeji Jun

**Writing – original draft:** Hye Lim Bae

**Writing – review & editing:** Kwangsoo Kim, Young Jun Chai

